# p63 co-opts the skin Krt8-to-Krt5 transition for enamel organ development

**DOI:** 10.1101/2025.02.11.637463

**Authors:** Qinghuang Tang, Jung-Mi Lee, Liwen Li, Chunmiao Cai, Hunmin Jung, Hyuk-Jae Edward Kwon

## Abstract

Tooth enamel, the hardest vertebrate tissue, is crucial for mastication and dental protection. Its formation depends on the enamel organ (EO), a specialized epithelial structure derived from oral epithelium. How uniform oral epithelium differentiates into diverse EO cell types remains unclear. While *p63*, an ectodermal development master regulator, is essential for dental placode formation, its specific roles in EO development have been obscured by early arrest in p63 knockout mice. Using single-cell RNA sequencing from mouse incisors, we show *p63* expression across all EO cell types with both shared and distinct functions. Through trajectory reconstruction, we identify *p63*’s role in regulating both amelogenic (AmG) and non-AmG lineage commitment during EO development. Comparative transcriptome analyses reveal that *p63* regulates the Krt8-to-Krt5 transition during AmG cell differentiation, paralleling its function in skin development. This parallel is reinforced by comparative motif discovery showing shared transcription factor usage, particularly p63 and AP-2 family members. Chromatin accessibility analyses further illustrate that *p63* mediates this transition through chromatin landscape remodeling. These findings demonstrate that *p63* co-opts the Krt8-to-Krt5 transition mechanism from skin development for EO formation.

**Summary statement:** This study reveals how p63 repurposes a skin keratinization mechanism for tooth enamel formation, providing novel insights into how specialized dental tissues develop and potential therapeutic targets for enamel disorders.

## Introduction

Tooth enamel, the hardest and most mineralized tissue in vertebrates, is essential for masticatory function and tooth protection (Lacruz et al., 2017). Enamel development is orchestrated by ameloblasts, specialized cells of the enamel organ (EO), which originate from the oral ectoderm (Liu et al., 2016). During EO development, the oral epithelium invades the dental mesenchyme and differentiates into four distinct cell types: inner enamel epithelium (IEE), stratum intermedium, stellate reticulum, and outer enamel epithelium (Smith and Nanci, 1995). The IEE subsequently differentiates into ameloblasts, which form the enamel matrix. While these major EO cell types are well characterized, the mechanisms governing oral epithelial specification into diverse EO cell lineages remain poorly understood.

The oral epithelium shares characteristics with other surface epithelia, comprising stratified squamous epithelium whose thickness and keratinization vary by location and functional requirements (Presland and Dale, 2000). Its development follows common stratification processes similar to other surface epithelia, including skin (Romano et al., 2012). During epithelial stratification, cells undergo morphological, biochemical, and transcriptional changes, with keratins serving as key markers that define cell identity and differentiation status (Roberts and Horsley, 2014; Cohen et al., 2022). In skin development, this stratification process involves a characteristic transition where single-layered surface ectoderm expressing Keratin 8 (Krt8) transitions to stratified epidermal progenitor cells through Krt8 downregulation and Krt5 upregulation (Blanpain and Fuchs, 2006).

The transcription factor p63, particularly its N-terminally truncated isoform Δ*Np63*, is a master regulator of stratified epithelial development (Soares and Zhou, 2018; Barbieri and Pietenpol, 2006). In skin, *p63* functions span from chromatin remodeling for epithelial lineage commitment (Yu et al., 2021; Li et al., 2019) to regulating terminal differentiation (Koster et al., 2007), including driving the Krt8-to-Krt5 transition through chromatin remodeling (Fan et al., 2018). Beyond the skin, *p63* is also crucial for tooth development, with Δ*Np63* widely expressed throughout all stages of EO development (Rufini et al., 2011). Mice lacking *p63* (*p63*^*−/−*^) exhibit arrested tooth development at the dental lamina stage (Laurikkala et al., 2006). Notably, patients with *p63*-related ectrodactyly-ectodermal dysplasia-clefting (EEC) syndrome often exhibit defective enamel deposition and dental anomalies (Rinne et al., 2006; Kantaputra et al., 2012; Koul et al., 2014), suggesting a regulatory role of *p63* in EO development. However, because *p63*^*−/−*^ mice fail to initiate tooth development, the specific role of *p63* during EO development remains unknown. Furthermore, whether the cellular state transitions observed in skin contribute to EO development, and how *p63* mediates these transitions, remains unexplored.

To address these knowledge gaps, we analyzed publicly available single-cell RNA sequencing (scRNA-seq) and chromatin accessibility datasets to investigate *p63*’s specific cellular and molecular roles in EO development. Our findings demonstrate that *p63* co-opts the Krt8-to-Krt5 cellular transition, previously established in skin development, to orchestrate epithelial differentiation during EO development. This study uncovers a conserved *p63*-driven regulatory mechanism for epithelial keratinization across different tissues, providing new insights into epithelial plasticity during development.

## Results

### p63 exerts both common and distinct functions in various cell types during enamel organ development

Genetic mutations in *p63* cause ectodermal dysplasia, a group of rare hereditary disorders affecting ectodermal organs including skin, sweat glands, and teeth (Romano et al., 2012). Patients with these mutations often exhibit various dental abnormalities, including tooth agenesis, alterations in tooth shape and size, fused teeth, and enamel dysplasia (Kantaputra et al., 2012). Despite this clear link between *p63* mutations and dental anomalies, *p63*’s specific role in enamel organ (EO) development and amelogenesis remains poorly understood. This knowledge gap exists largely because germline *p63* knockout mice exhibit tooth developmental arrest at the lamina stage, before the onset of amelogenesis (Laurikkala et al., 2006). The advent of scRNA-seq technique has revolutionized our ability to examine gene expressions at unprecedented resolution (Jovic et al., 2022). To understand p63’s functions in EO development, we revisited a scRNA-seq dataset from continuously growing adult mouse incisor teeth, which contain all stages of EO cell differentiation from stem/progenitors to terminally differentiated ameloblasts (Sharir et al., 2019). Using Seurat (Stuart et al., 2019), we clustered EO cells into 15 subtypes and assigned putative cell identities based on their distinct gene expression signatures (Fig. 1A). As previously reported (Sharir et al., 2019), these 15 EO cell types formed a transcriptomic continuum in UMAP space and could be grouped into three classes, with ‘cycling cells’ giving rise to both amelogenic (AmG) and non-AmG EO cells (Fig. 1A, red dashed arrows). While *p63* was broadly expressed across all 15 EO cell types (Fig. 1B), consistent with previous *in situ* hybridization results (Laurikkala et al., 2006), single-cell resolution revealed that not every individual cell expressed *p63* (Fig. 1B). Notably, *p63* showed highest expression in the inner stellate reticulum/stratum intermedium (ISR/SI) cells (Fig. 1C), a subset of non-AmG EO cells that serve as a stem cell reservoir for ameloblast renewal (Liu et al., 2016). This finding suggests a possible role for *p63* in dental stem/progenitor cell replenishment. To further elucidate *p63*’s specific roles in different EO cell types, we identified the differentially expressed genes (DEGs) between *p63* positive (*p63*^+^) and *p63* negative (*p63*^*−*^) cells across all EO cell types. Functional classification of these DEGs revealed that *p63* exerts distinct functions in different EO cell types, while maintaining shared roles in p53 signal transduction, chromatin remodeling and odontogenesis (Fig. 1D). Together, these results demonstrate *p63*’s essential and diverse roles in regulating EO development.

**Figure 1.**
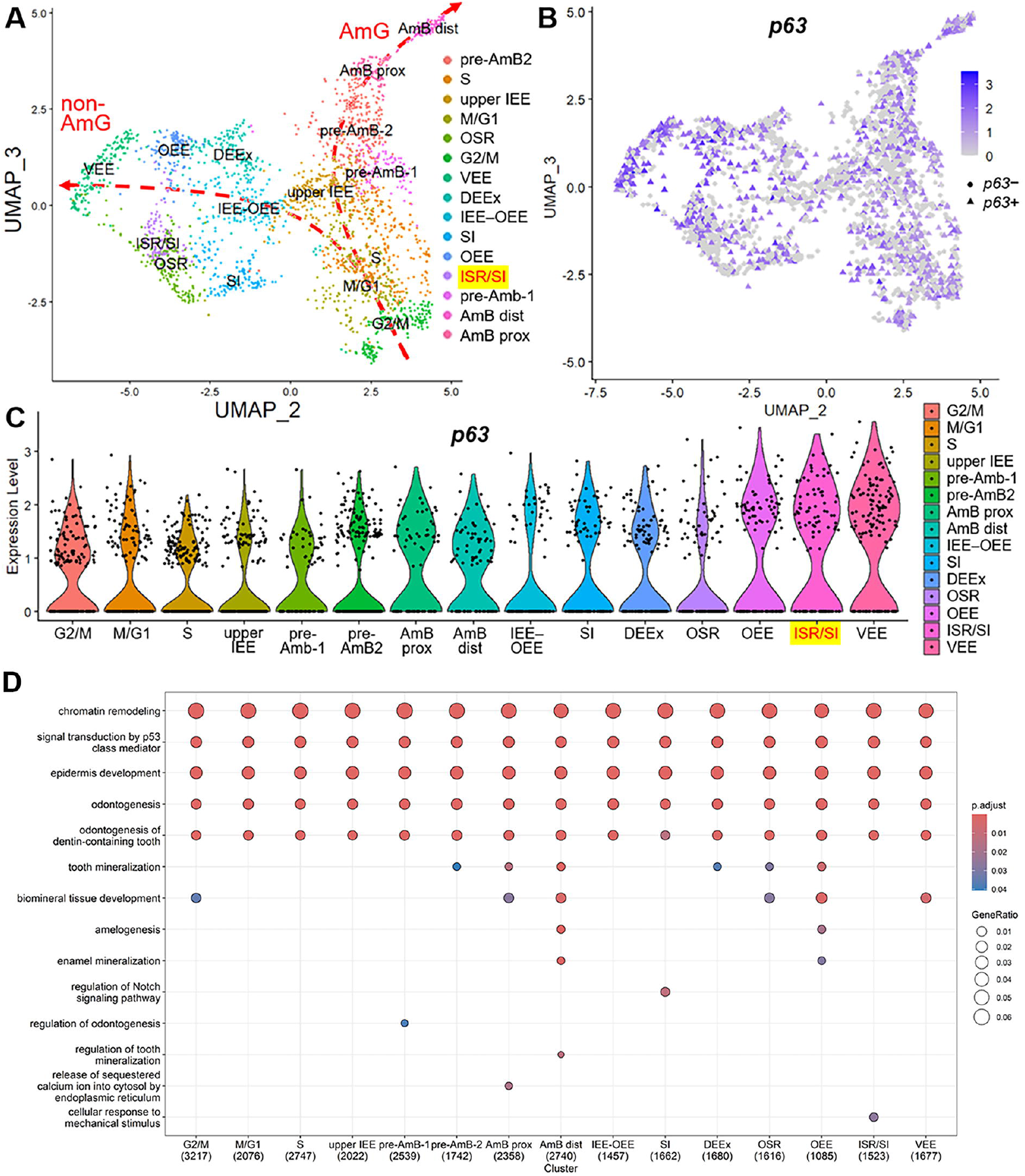
*p63* exhibits diverse functional roles during enamel organ development. (**A**) DimPlot illustrating major dental epithelial cell types in the 8-week-old mouse incisor. (**B**) Dynamic distributions of *p63*-expressing (*p63*^+^) cells across major dental epithelial cell types. (**C**) VlnPlot demonstrating different expression levels of *p63* in major dental epithelial cell types. (**D**) Gene ontology (GO) analyses of differentially expressed genes (DEGs) between *p63*^+^ vs *p63*^-^ groups in each dental epithelial cell type. upper IEE, upper inner enamel epithelium; OEE, outer enamel epithelium; VEE, ventral enamel epithelium; DEEx, dental epithelial extensions; IEE– OEE, junction between the IEE and OEE; SI, stratum intermedium; ISR, inner stellate reticulum; OSR, outer stellate reticulum; pre-AmB, pre-ameloblasts; AmB prox, proximal ameloblasts; AmB dist, distal ameloblasts; G2/M, M/G1 and S, cycling dental epithelial cells.

### *p63* participates in both AmG and non-AmG lineage commitment during EO development

Given *p63*’s crucial role in epidermal lineage commitment (Li et al., 2019), we investigated its functions in EO cell fate specification by reconstructing the developmental trajectory of EO cells using Monocle2 (Qiu et al., 2017). The analysis revealed a branched continuum with four branch points representing cellular fate decisions. At branch point 1 (Fig. 2A, circle 1), cycling cells bifurcated into AmG and non-AmG lineages. Notably, *p63* showed pseudo-temporal expression in both AmG and non-AmG processes during EO cell fate specification (Fig. 2A). Analysis of genes with branch point-dependent expression revealed that *p63*, along with other EO stem cell marker genes (Gan et al., 2020; Yu et al., 2020), functions as a branching gene at all four branch points (Fig. S1A–D). We then focused specifically on the branching genes responsible for AmG and non-AmG lineage bifurcation at branch point 1 (Fig. 2B). Gene ontology (GO) analysis of these genes demonstrated enrichment for epidermal cell-related biological processes, including odontogenesis (Fig. 2C). Importantly, *p63* was found to be involved in odontogenesis-related biological processes (Fig. 2D). Together, these results suggest that *p63* functions as a potential regulator in the temporal specification of both AmG and non-AmG EO cell fates during EO development, with particular importance in the bifurcation of AmG and non-AmG lineages (Fig. S1E).

**Figure 2.**
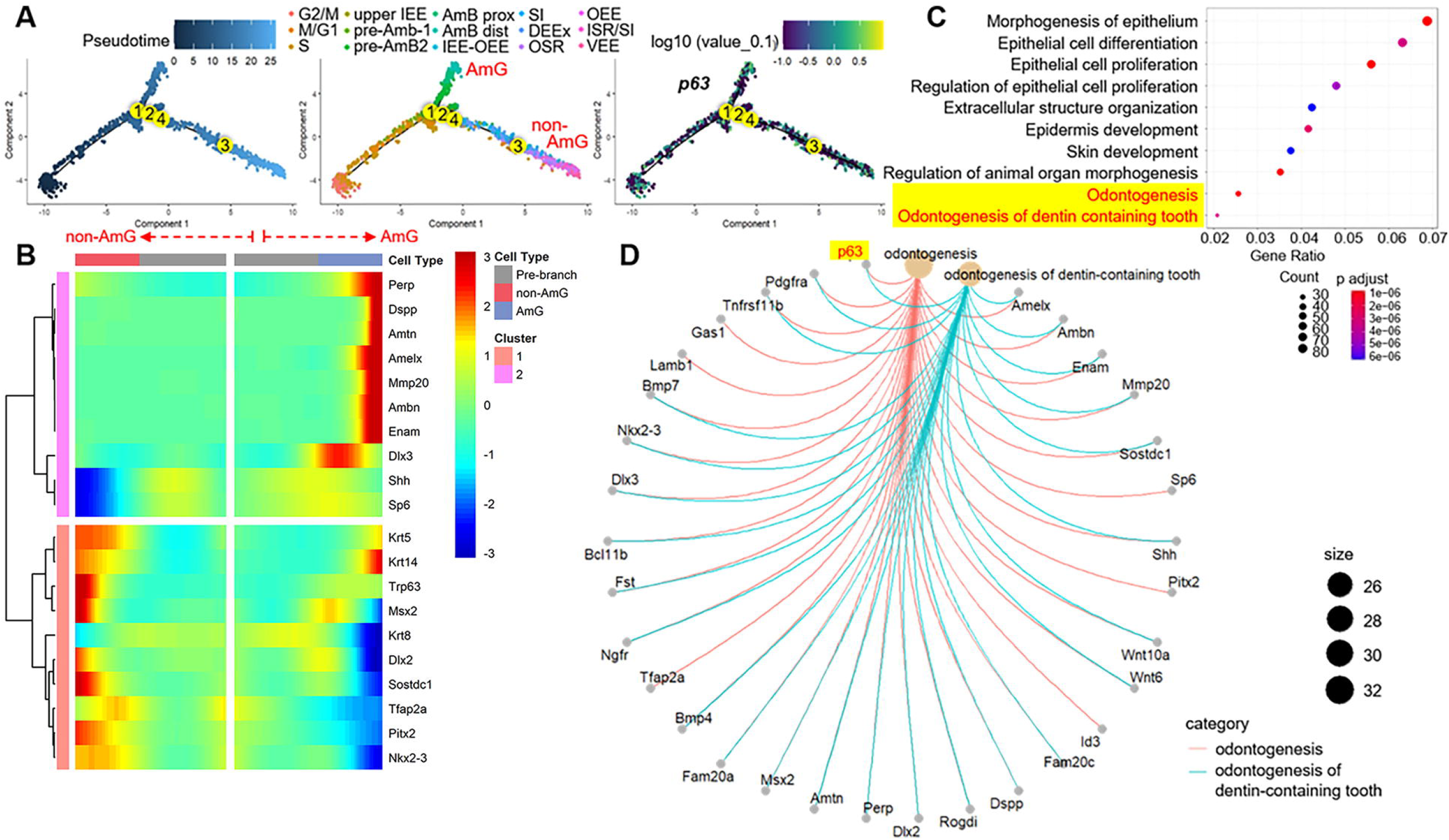
*p63* is a regulator of lineage bifurcation in enamel organ development. (**A**) Pseudotime analysis illustrating the divergence of amelogenic (AmG) and non-AmG lineages within dental epithelial cells. As per the updated model, a population of actively cycling progenitors give rise to both AmG and non-AmG epithelial cells during dental epithelial cell specification. Notably, *p63* is consistently expressed throughout the entire pseudotime trajectory. The numbers 1–4 indicate the branching points along the pseudotime trajectory. (**B**) Branched heatmap depicting the expression patterns of representative branching genes, including *p63*, during the bifurcation of AmG and non-AmG epithelial cells at branching point 1. (**C**) GO analysis of the branching genes identified at branching point 1. (**D**) cnetplot demonstrating that p63 is a key player in odontogenesis-related biological processes.

### Krt8-to-Krt5 cellular state transition occurs in both AmG and non-AmG cells during EO development

The oral cavity’s epithelial layer consists of stratified squamous epithelium with varying degrees of stratification and keratinization, reflecting the functional demands of specific anatomical regions (Presland and Dale, 2000; Romano et al., 2012). Among epithelial-derived structures, the enamel organ is unique in its ability to form enamel, the hardest vertebrate tissue. Previous studies in mammalian skin have established *p63*’s genome-wide effect on epidermal cell fate commitment (Li et al., 2019), particularly in governing the transformation of Krt8^+^ progenitor cells to Krt5^+^ epidermal progenitors, known as the Krt8-to-Krt5 transition (Fan et al., 2018). To investigate whether this *p63*-directed epidermal lineage formation mechanism applies to EO cell lineage specification, we examined *Krt8* and *Krt5* expression in EO cells (Fig. 3A, B). *Krt8* expression was higher at the root of the EO cell transcriptomic continuum (Fig. 3A, arrow-tail), while *Krt5* expression was elevated at the differentiation trajectory’s end (Fig. 3B, arrowheads). Gene expression analysis along pseudotime revealed decreasing *Krt8* expression (Fig. 3E) and increasing *Krt5* expression (Fig. 3F) during EO cell differentiation, indicating a Krt8-to-Krt5 transition. This Krt8-to-Krt5 transition was observed in both AmG (Fig. 3C) and non-AmG EO cell differentiation (Fig. 3G, H), as evidenced by the inverse expression trends of *Krt8* and *Krt5* along each cell lineage (Fig. 3C, G, H). Quantification analyses of expression percentage (Fig. S2A) and levels (Fig. S2B) of *Krt8, Krt5* and *p63* showed that *p63* expression coincides with the Krt8-to-Krt5 transition, evidenced by p63’s positive correlation with Krt5 and negative correlation with Krt8 in both cell lineages. Notably, along the AmG cell differentiation trajectory, *p63* expression peaked at an intermediate stage during the Krt8-to-Krt5 transition (Fig. 3D). These results indicate that p63 may play a role in regulating the Krt8-to-Krt5 transition during EO development, particularly during AmG cell differentiation.

**Figure 3.**
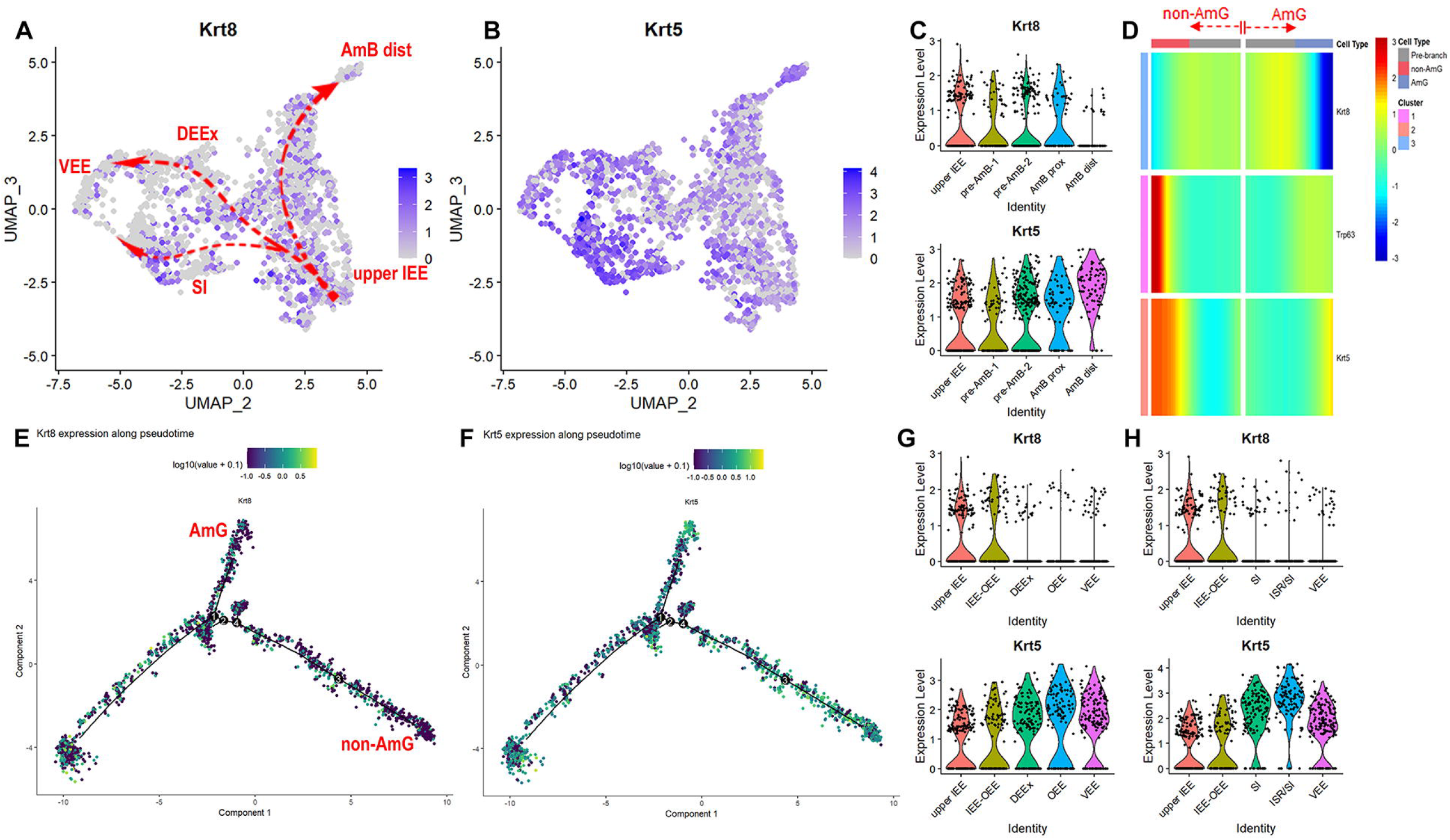
The occurrence of Krt8-to-Krt5 cellular state transition during enamel organ development. **(A, B**) FeaturePlots showing the expressions of *Krt8* (A) and *Krt5* (B) in major dental epithelial cell types. (**C**) VlnPlot depicting dynamic expression changes of *Krt8* and *Krt5* during AmG lineage cell specification. (**D**) Branched heatmap showing *p63* expression during the Krt8-to-Krt5 transition at the AmG and non-AmG bifurcation point. (**E, F**) Pseodotemporal expression of *Krt8* (E) and *Krt5* (F) during the AmG and non-AmG bifurcation. (**G, H**) VlnPlot showing dynamic expression changes of *Krt8* and *Krt5* during non-AmG lineage cell specification. upper IEE, upper inner enamel epithelium; DEEx, dental epithelial extensions; SI, stratum intermedium; VEE, ventral enamel epithelium; AmB dist, distal ameloblasts.

### *p63* regulates Krt8-to-Krt5 transition in AmG Cells during EO development

To examine *p63*’s role in regulating the Krt8-to-Krt5 transition during AmG cell differentiation, we conducted a detailed analysis of the AmG cell lineage. During AmG cell differentiation, *p63* expression in major AmG cell types showed a gradual increase, accompanied by decreasing *Krt8* and increasing *Krt5* expression (Fig. 4A), characteristic of a Krt8-to-Krt5 transition. AmG cell trajectory analysis confirmed that *p63* expression pseudo-temporally coincides with this Krt8-to-Krt5 transition (Fig. 4B). To further investigate *p63*’s regulatory role, we performed differential gene expression analysis comparing two groups in the AmG cells: *p63*^+^ vs. *p63*^*−*^ (p63-DEGs) and *Krt5*^+^ vs. *Krt8*^+^ (Krt-DEGs). We identified 4,935 p63-DEGs and 4,987 Krt-DEGs, with substantial overlap of 4,753 shared DEGs (Fig. 4C). Analysis of transcriptomic changes between these groups, based on DEG ranking by fold change and p-value, revealed similar patterns (Fig. 4D). GO analysis of Krt-DEGs showed significant enrichment in ‘signal transduction by p53 class mediator’ (Fig. 4E). These findings suggest that *p63* regulates the Krt8-to-Krt5 transition during AmG cell differentiation.

**Figure 4.**
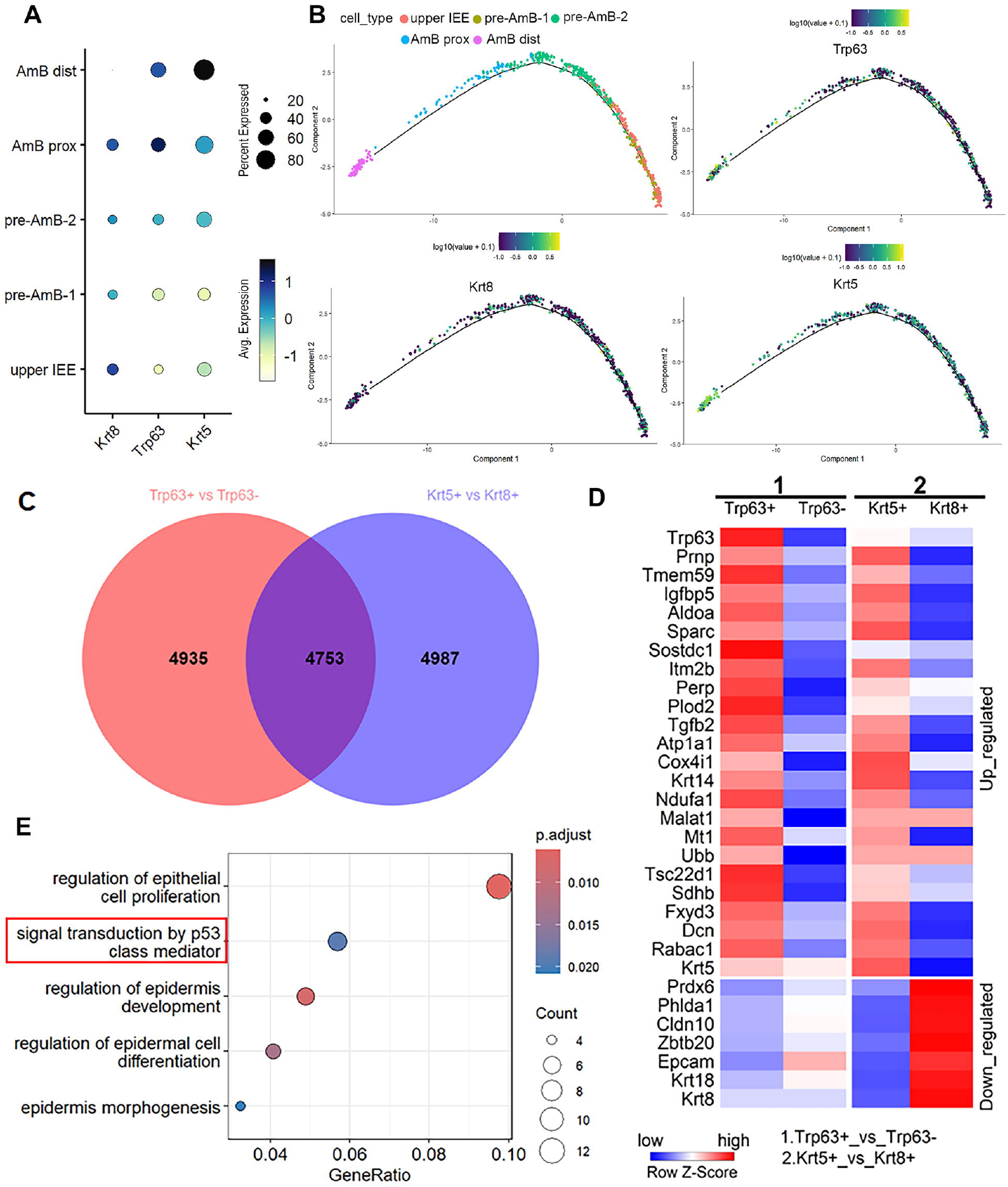
*p63* regulates the Krt8-to-Krt5 transition during AmG cell differentiation. **(A)** Dot Plots showing the concurrent expression of *p63* with *Krt5* during the amelogenic Krt8-to-Krt5 transition. (**B**) Pseudotemporal expression changes of *Krt8, Krt5* and *p63* throughout amelogenesis. (**C**) Venn diagram illustrating the significant overlap (4753 genes) of differential expressed genes (DEGs) between the *p63*^+^ vs. *p63*^*−*^ and *Krt5*^+^ vs. *Krt8*^+^ dental epithelial cell groups. (**D**) Heatmap displaying the similarity of transcriptomic changes between group 1 (*p63*^+^ vs. *p63*^*−*^) and group 2 (*Krt5*^+^ vs. *Krt8*^+^). (**E**) GO analysis of DEGs in the *Krt5*^+^ vs. *Krt8*^+^ group.

### *p63* regulates Krt8-to-Krt5 transition during AmG cell differentiation through chromatin remodeling

Previous research showed that *p63*-governed chromatin accessibility changes drive the Krt8-to-Krt5 transition in the skin (Fan et al., 2018). To investigate whether *p63* similarly regulates this transition during AmG cell differentiation, we analyzed the chromatin accessibility of *Krt8* and *Krt5* in incisor development at embryonic day 12 (E12) and E16 (Wang et al., 2022). Comparative analyses revealed opposing dynamics: the *Krt8* locus showed decreased chromatin accessibility as incisor development progressed (Fig. S3A), while the *Krt5* locus showed increased accessibility (Fig. S3B), indicating that chromatin remodeling also underlies the Krt8-to-Krt5 transition during AmG cell differentiation. To investigate *p63*’s role in this process, we performed comparative motif analyses on the cis-regulatory elements of Krt-DEGs and on regions that gained chromatin accessibility during the skin Krt8-to-Krt5 transition (Fig. 5A). *De novo* motif discovery using HOMER (Heinz et al., 2010) identified p63 and TFAP2C as the two most enriched transcription factors (TFs) in regions gaining chromatin accessibility during skin transition (Fig. 5C). Interestingly, motif scan using BETA (Wang et al., 2013) identified AP-2 family and p53 class TFs as the most enriched TF groups in Krt-DEG cis-regulatory elements (Fig. 5B), suggesting conserved AP-2 and p53 family TF usage during Krt8-to-Krt5 transition in both AmG and skin cells. Furthermore, gene set enrichment analysis of ranked p63-DEGs from AmG cells showed strong enrichment for both genes associated with regions gaining chromatin accessibility during skin Krt8-to-Krt5 transition (Fig. 5D) and *Krt5*^+^ AmG cell signature gene set (Fig. 5E). More importantly, *p63* knockout (KO) inversely altered chromatin accessibility dynamics at the candidate cis-regulatory elements (cCREs) of *Krt8* and *Krt5* in the skin (Fig. 5F). Notably, these ENCODE-compiled cCREs contained p63 binding sites (Fig. 5F) that overlap with differentially accessible regions in incisor (Fig. S3A, B). These results indicate that *p63* regulates the Krt8-to-Krt5 transition through chromatin landscape remodeling during AmG cell differentiation in the EO. Collectively, our study provides convincing evidence that *p63* co-opts the prevalent Krt8-to-Krt5 transition in skin for EO development.

**Figure 5.**
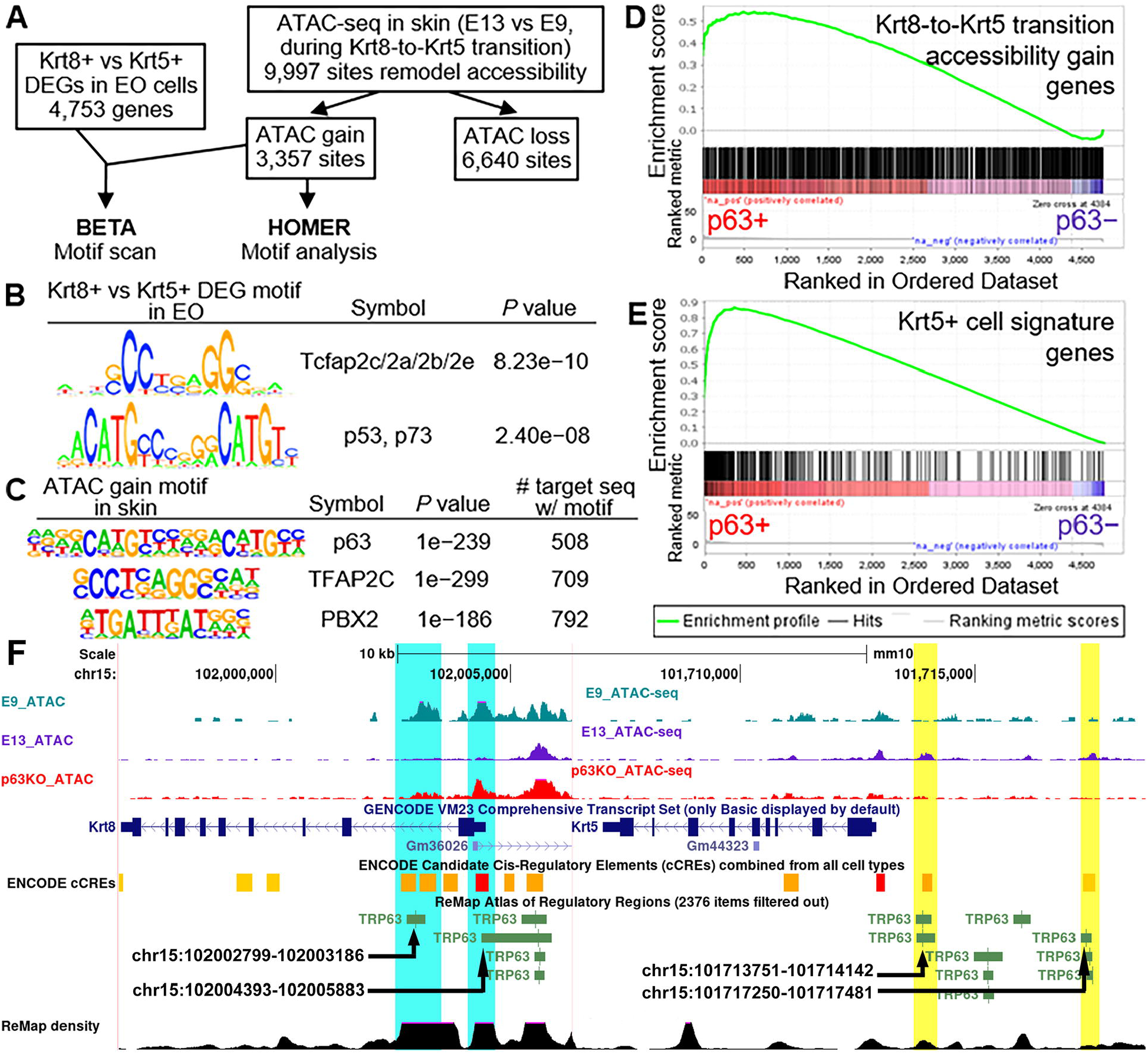
*p63* remodels open chromatin regions during the Krt8-to-Krt5 transition in AmG cell differentiation. **(A)** Schematic representation of the integrative analysis workflow comparing enriched motifs in *Krt8*^+^ vs. *Krt5*^+^ DEGs from dental epithelium and genomic loci (E13 vs. E9 ATAC-seq differential binding sites) that gained chromatin accessibility during the Krt8-to-Krt5 transition in skin. (**B**) Motif scan analysis revealing high enrichment of p53 and AP-2 family transcription factors (TFs) in DEGs between *Krt8*^+^ vs. *Krt5*^+^ dental epithelial cells. (**C**) Motif enrichment analysis demonstrating high enrichment of p63 and AP-2 family TFs in gained open chromatin regions during the Krt8-to-Krt5 transition in skin. (**D, E**) Gene set enrichment analysis (GSEA) of ranked DEGs between *p63*^+^ vs. *p63*^*−*^ dental epithelial cells against (D) genes associated with gained chromatin regions during the Krt8-to-Krt5 transition in skin and (E) signature genes of *Krt5*^+^ dental epithelial cells. (**F**) ATAC-seq tracks revealing that *p63* knockout reverses chromatin accessibility dynamics in the candidate cis-regulatory elements (cCREs) of *Krt8* (turquoise) and *Krt5* (yellow). cCREs are combined from all cell types by ENCODE. Multiple p63 binding sites are present in the cCREs of *Krt8* and *Krt5*, as visualized by the ReMap track. TRP63 binding sites with labeled chromosome coordinates indicate their presence in differentially accessible regions during incisor development.

## Discussion

Our study demonstrates that *p63* co-opts the Krt8-to-Krt5 transition, previously established in skin development, to orchestrate EO development. Through scRNA-seq analysis, we show that *p63* is expressed across all EO cell types and participates in both AmG and non-AmG lineage commitment. Importantly, we identify that *p63* regulates the Krt8-to-Krt5 transition during AmG cell differentiation through chromatin landscape remodeling, similar to its role in skin development.

### p63 is a ubiquitous player during EO development

While major EO cell types are well documented (Smith and Nanci, 1995), the mechanisms controlling their specification remain incompletely understood. Understanding these mechanisms is crucial since precise and proper EO cell coordination ensures functional enamel formation (Lacruz et al., 2017). As a master regulator of ectodermal development, p63 integrates key signaling pathways in early tooth development for dental placode formation (Laurikkala et al., 2006), but early dental arrest in p63^*−*/*−*^ mice has limited our understanding of its later roles in EO development. Our scRNA-seq data reanalysis unravels that p63 regulates shared functions across EO cell types, including p53 signaling, chromatin remodeling, and odontogenesis, while also controlling cell-type-specific processes like amelogenesis and mineralization (Fig. 1). Trajectory analysis reveals that p63 participates in both AmG (Fig. 2) and non-AmG (Fig. S2) lineage commitment during EO development, acting as a branching gene. These results indicate that p63 orchestrates EO development by controlling EO cell lineage commitment and differentiation, aligning with previous study demonstrating p63 expression in both undifferentiated EO cells and differentiated ameloblasts (Rufini et al., 2006). In the current study, we identified DEGs between *p63*^+^ and *p63*^*−*^ cells across different EO cell types to infer *p63*’s role in EO development (Fig. 1, 4 and 5). Given that enamel formation requires precise coordination between different EO cell types (Liu et al., 2016), future studies investigating cell-cell communications between these different cell populations would provide additional insights into how *p63* orchestrates EO development through potential paracrine signaling mechanisms.

### *p63* orchestrates a conserved regulatory mechanism for epithelial keratinization in EO development

Surface epithelia, including oral epithelium, form essential protective barriers against chemical, microbial, and physical challenges, making them indispensable for organismal viability (Presland and Dale, 2000). To establish and maintain this protective function, surface epithelial cells undergo a tightly regulated differentiation program that results in the formation of structural proteins, particularly keratins (Roberts and Horsley, 2014). These keratins serve as critical markers of cellular state transitions in epithelial biology, playing essential roles in maintaining tissue homeostasis and adapting to developmental cues (Cohen et al., 2022). Recent studies have revealed that mutations and genetic variants in keratins increase dental decay risk and susceptibility (Duverger et al., 2018; Duverger et al., 2014), highlighting the importance of understanding keratin regulation in EO development. Through reanalysis of incisor scRNA-seq data, we discovered that the Krt8-to-Krt5 transition, a hallmark of epidermal fate commitment in the skin (Fan et al., 2018), occurs during EO cell differentiation (Fig. 3). The AmG Krt8-to-Krt5 transition coincided with pseudo-temporal *p63* expression (Fig. 4A, B), consistent with previous study showing that p63 colocalizes with Krt5 during EO development (Rufini et al., 2006). Similar transcriptome changes between *p63*^+^ vs *p63*^*−*^ and *Krt5*^+^ vs *Krt8*^+^ EO cells suggested *p63* regulation of the Krt8-to-Krt5 transition during AmG cell differentiation (Fig. 4C-E). Supporting this, gene set enrichment analyses of DEGs between *p63*^+^ vs *p63*^*−*^ cells showed strong enrichment of *Krt5*^+^ cell signature genes (Fig. 5D, E). Motif analyses of chromatin remodeling regions for *Krt8* and *Krt5* during both skin (Fig. 5F) and incisor (Fig. S3) development both revealed the presence of p63 binding sites. Together, these findings demonstrate that *p63* co-opts the Krt8-to-Krt5 transition mechanism from skin development for EO development through chromatin landscape remodeling, revealing a conserved *p63*-driven regulatory mechanism for epithelial keratinization across different tissues. This finding is particularly significant as it establishes a molecular link between keratin regulation and enamel formation, potentially offering new therapeutic targets for enamel disorders.

Our multi-omics data re-analysis approach, combining single-cell transcriptomics and chromatin accessibility data, provides a powerful framework for understanding gene regulatory functions in contexts where traditional genetic approaches are limited by developmental arrest. Together, these findings not only reveal a conserved developmental mechanism across different epithelial tissues but also provide crucial insights into the molecular basis of enamel formation, opening new avenues for therapeutic interventions in enamel disorders.

## Materials and methods

### Analysis of scRNA-Seq data from mouse incisor

We analyzed a scRNA-seq dataset from adult mouse incisors (8–12 weeks, GSE131204) using the Seurat pipeline (Stuart et al., 2019). Quality control filtering (minimum 5 cells per gene, nFeature_RNA > 1,000, percent mitochondrial reads < 10%) yielded 3,171 high-quality cells. For enamel organ (EO) heterogeneity analysis, we excluded immune and mesenchymal cells based on established markers, retaining 2,341 EO cells. Following Seurat’s standard workflow, we identified 2,000 highly variable genes after log normalization and performed principal component analysis. The top 10 principal components, selected using the ElbowPlot function, were used for downstream analysis. We constructed a shared nearest-neighbor graph using the FindNeighbors function and identified 15 main cell clusters (resolution = 1.2) using the FindClusters function. Cell types were classified using cluster-specific marker genes from previous research (Sharir et al., 2019) and visualized using Uniform Manifold Approximation and Projection (UMAP). For *p63* expression analysis, cells with non-zero *p63* counts were classified as *p63*-positive (*p63*^+^). We identified differentially expressed genes (DEGs) between *p63*^+^ and *p63*-negative cells (p63-DEGs) using the FindMarkers function. Similarly, we determined DEGs between *Krt8*-positive and *Krt5*-positive cells (Krt-DEGs). These DEGs were ranked by combining p-value and average log fold change, then analyzed using gene set enrichment analysis (GSEA) software (Subramanian et al., 2005). To analyze EO cell differentiation trajectories, we employed Monocle2 (Qiu et al., 2017). Cell embeddings and clusters from Seurat were converted into a CellDataSet object and ordered in pseudotime. We assigned the trajectory root to the node containing the majority of actively cycling cells. Branch-dependent expression analysis was performed using the BEAM function (q-value < 0.01). Gene ontology analysis was conducted using the clusterProfiler R package (Yu et al., 2012).

### Analysis of ATAC-Seq data from mouse incisor and skin

We analyzed Assay for Transposase-Accessible Chromatin with sequencing (ATAC-seq) datasets from skin (E9 and E13, GSE97213) and incisor (E12 and E16, PRJNA668198) to investigate chromatin accessibility during the Krt8-to-Krt5 transition. For skin datasets, reads were aligned to the mouse mm10 reference genome using Bowtie2 (--local option) (Langmead and Salzberg, 2012) and processed with SAMtools (Li et al., 2009). After excluding mitochondrial and unplaced/random contigs, peaks were called using MACS2 (Zhang et al., 2008). For visualization in the UCSC genome browser, filtered BAM files were converted to bigwig format and merged by developmental stage. Differentially accessible chromatin regions between E13 and E9 were identified using the DiffBind R package (Ross-Innes et al., 2012). Regions showing increased accessibility during the skin Krt8-to-Krt5 transition underwent motif enrichment analysis using HOMER (Heinz et al., 2010), followed by motif scanning of Krt-DEGs cis-regulatory elements with BETA (Wang et al., 2013). Incisor ATAC-seq data was processed using the ENCODE pipeline (Hitz et al., 2023). Merged BAM files from each developmental stage were converted to bigWig format using the deepTools2 package (Ramirez et al., 2016) for IGV visualization. Accessibility differences between E16 and E12 were calculated using IGV’s subtract operation (Robinson et al., 2011).

## Acknowledgements

We thank Zhaoming Wu, PhD (The University of Hong Kong) for technical assistance with differential gene expression analysis and ranking of p63-DEGs and Krt-DEGs. We thank members of the Kwon laboratory for their insightful discussions and constructive feedback on this manuscript.

## Author contributions

Q. Tang, contributed to conception, design, data acquisition and interpretation, performed statistical analyses, drafted, and critically revised the manuscript. J.-M. Lee, L. Li, C. Cai, and H. Jung, contributed to data interpretation, and critically revised the manuscript. H.-J.E. Kwon, contributed to conception, design, and critically revised the manuscript. All authors gave final approval and agreed to be accountable for all aspects of the work.

## Data and resource availability

All relevant data and information regarding resources are available in the article and its supplementary information.

---

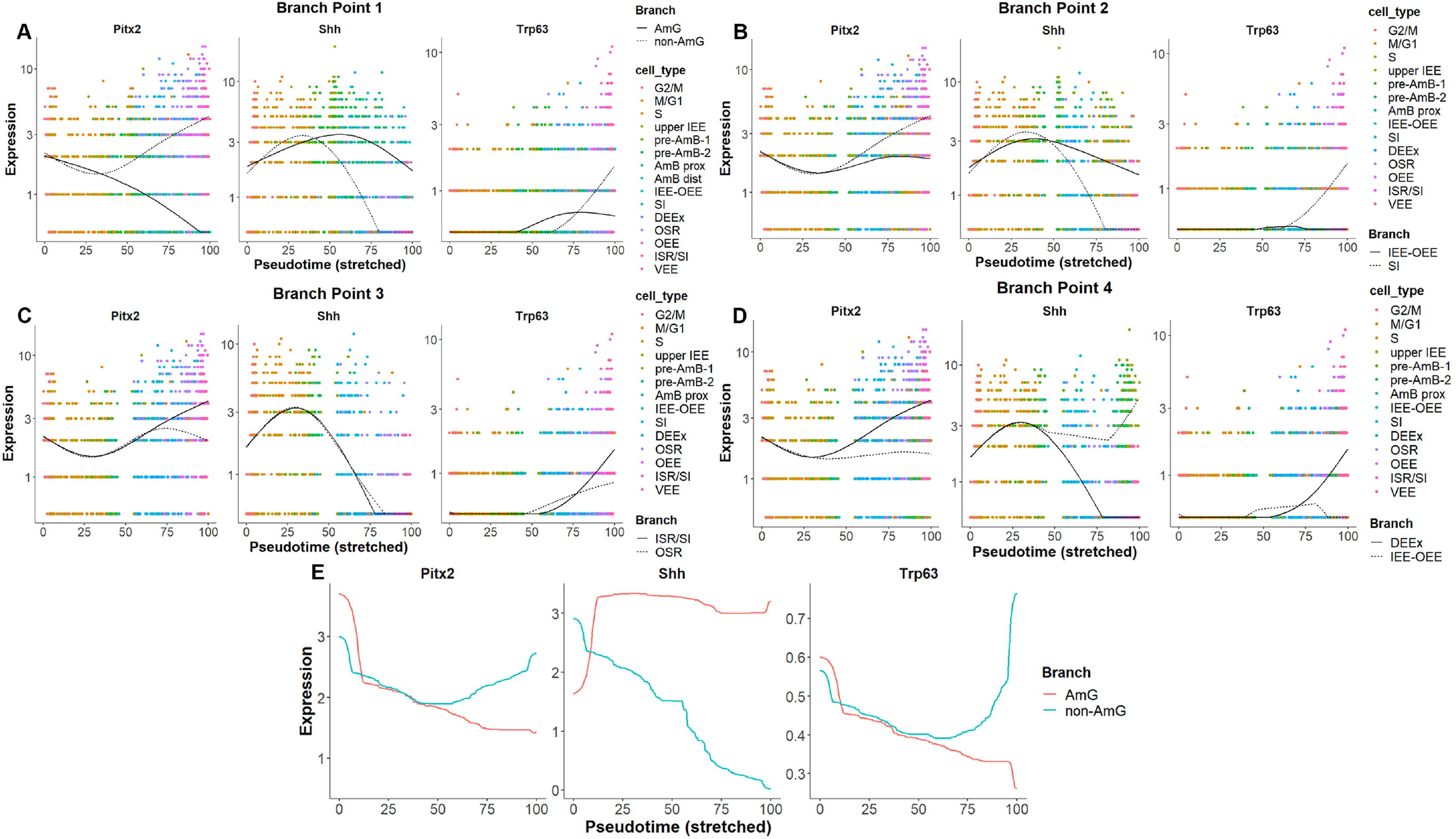

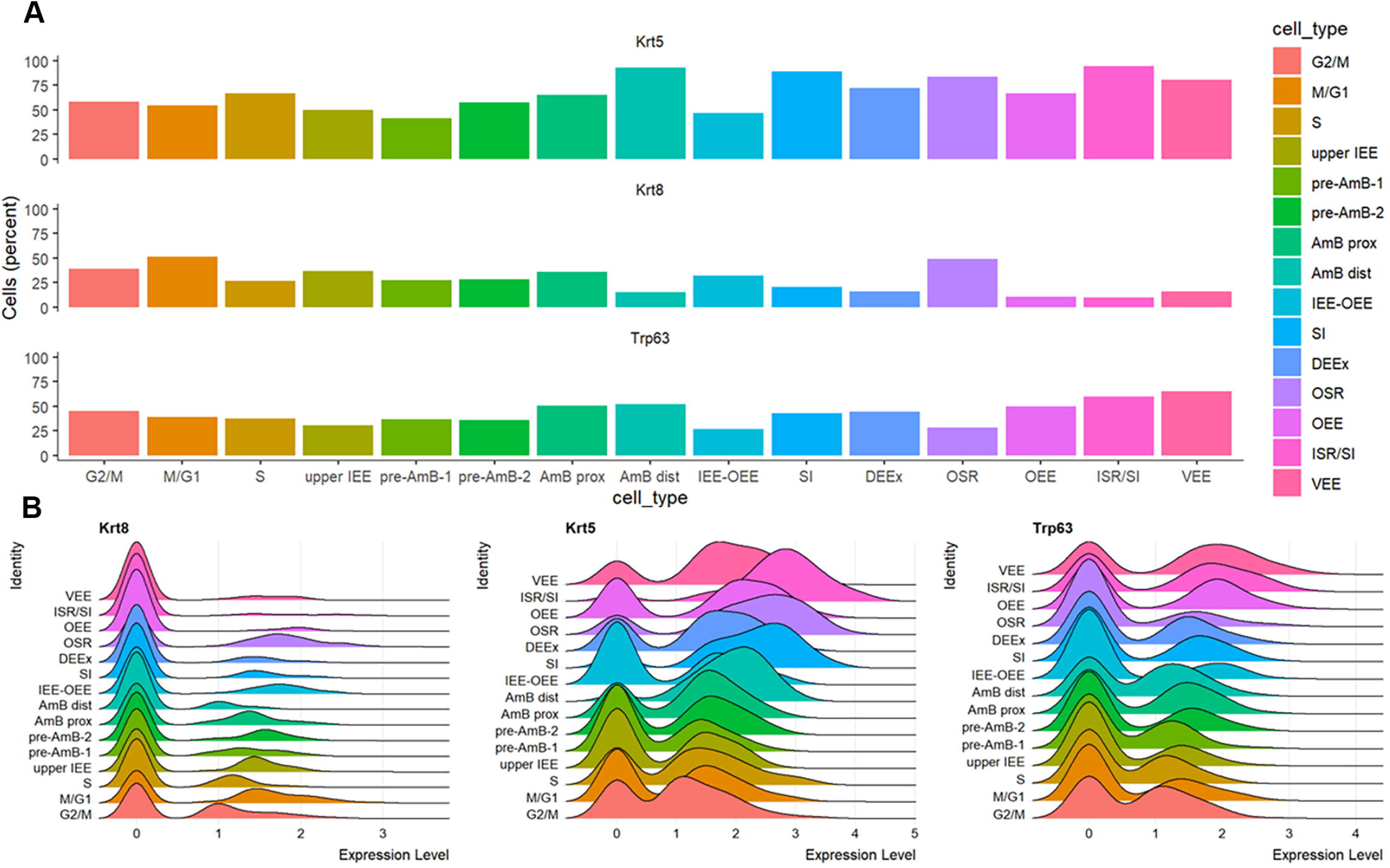

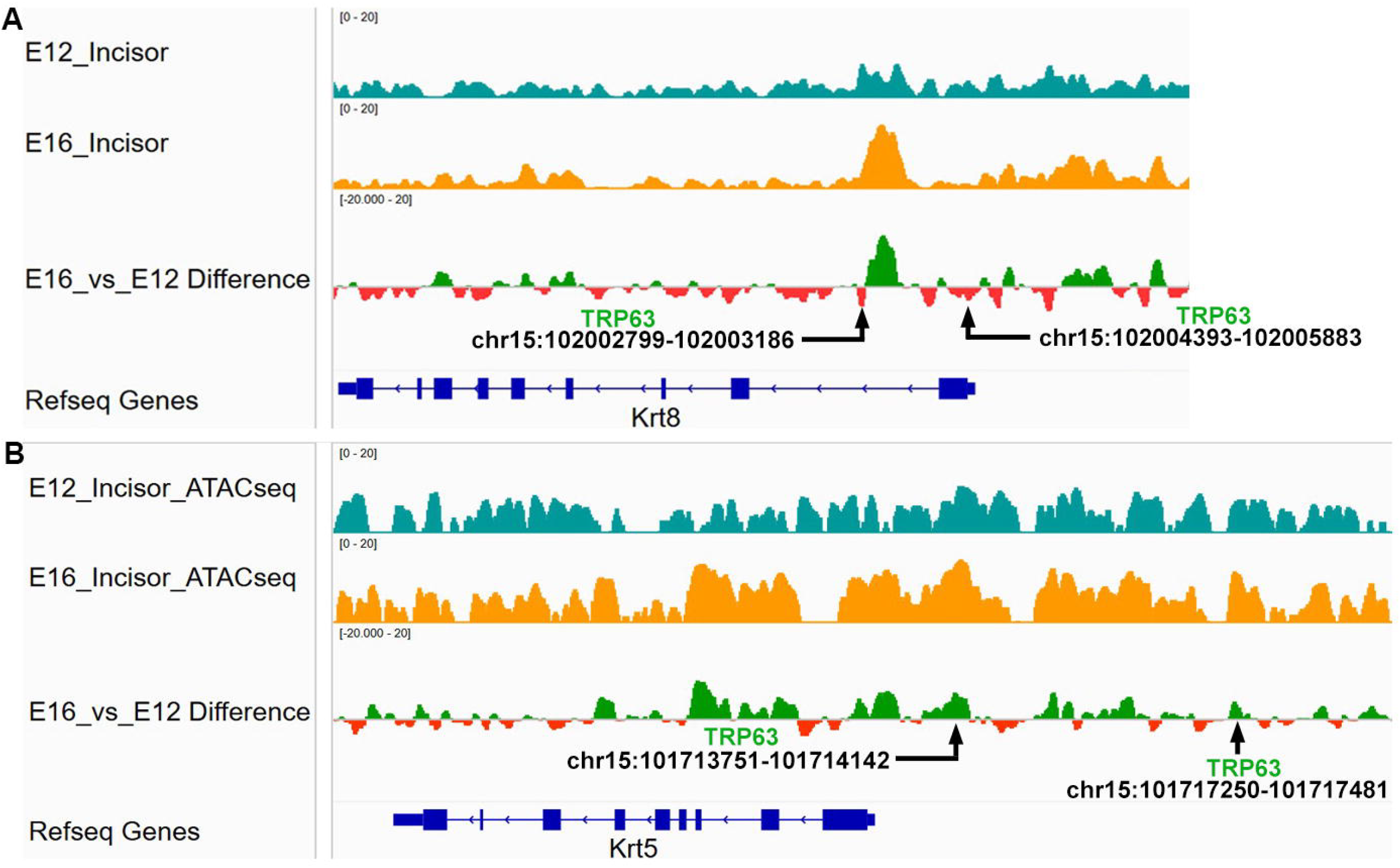

