## Supplemental information for "p63 co-opts the skin Krt8-to-Krt5 transition for enamel organ development"

**Supplementary information**

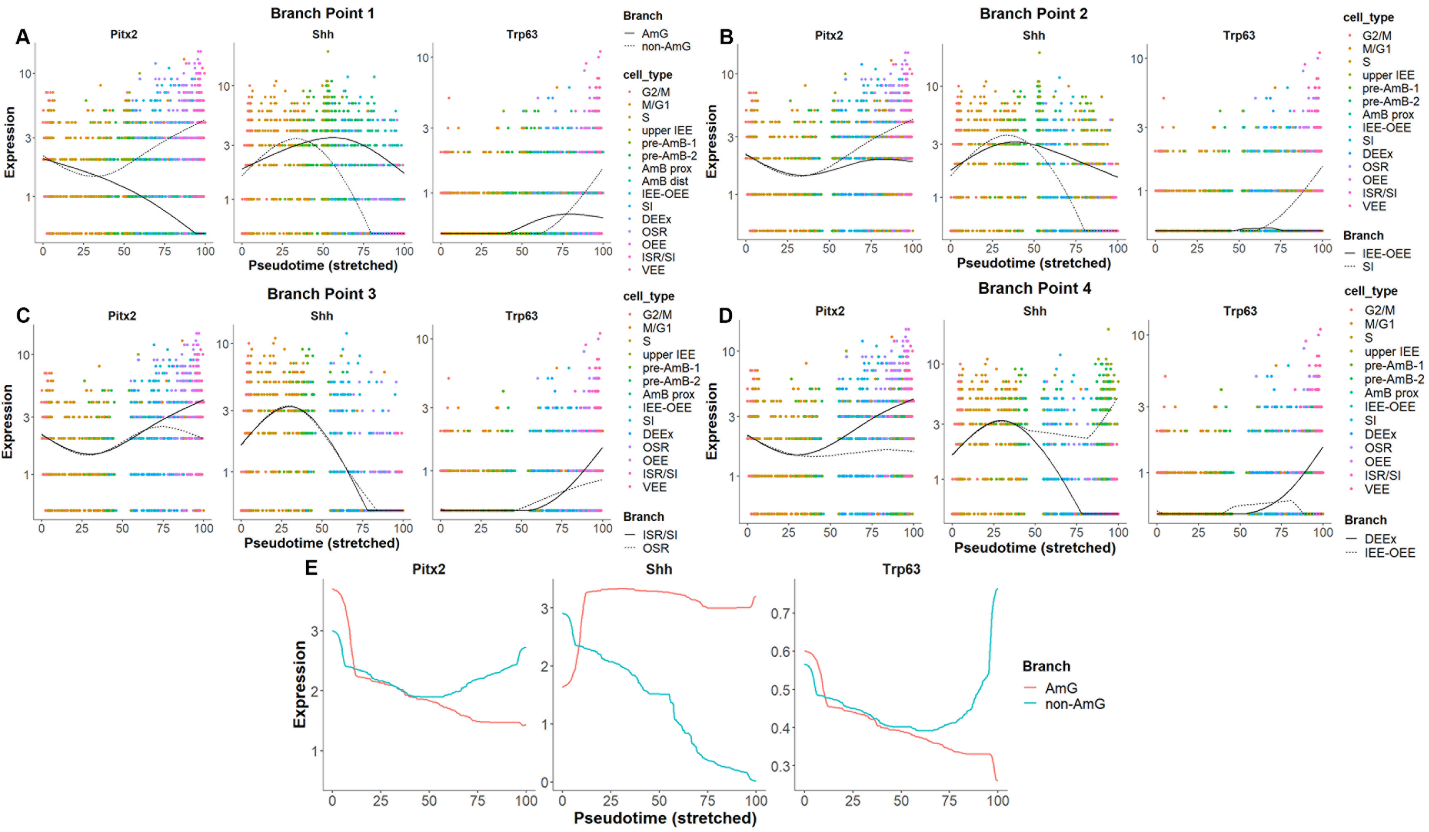

**Supplementary figure S1. *p63* functions as a branching gene throughout enamel organ development.** (**A–D**) *p63* acts as a branching gene at multiple branching points along the pseudotime trajectory of dental epithelial cell differentiation. (**E**) Kinetic curve along the pseudotime trajectory of dental epithelial cell differentiation demonstrating that *p63* is a regulator of lineage bifurcation during enamel organ development.

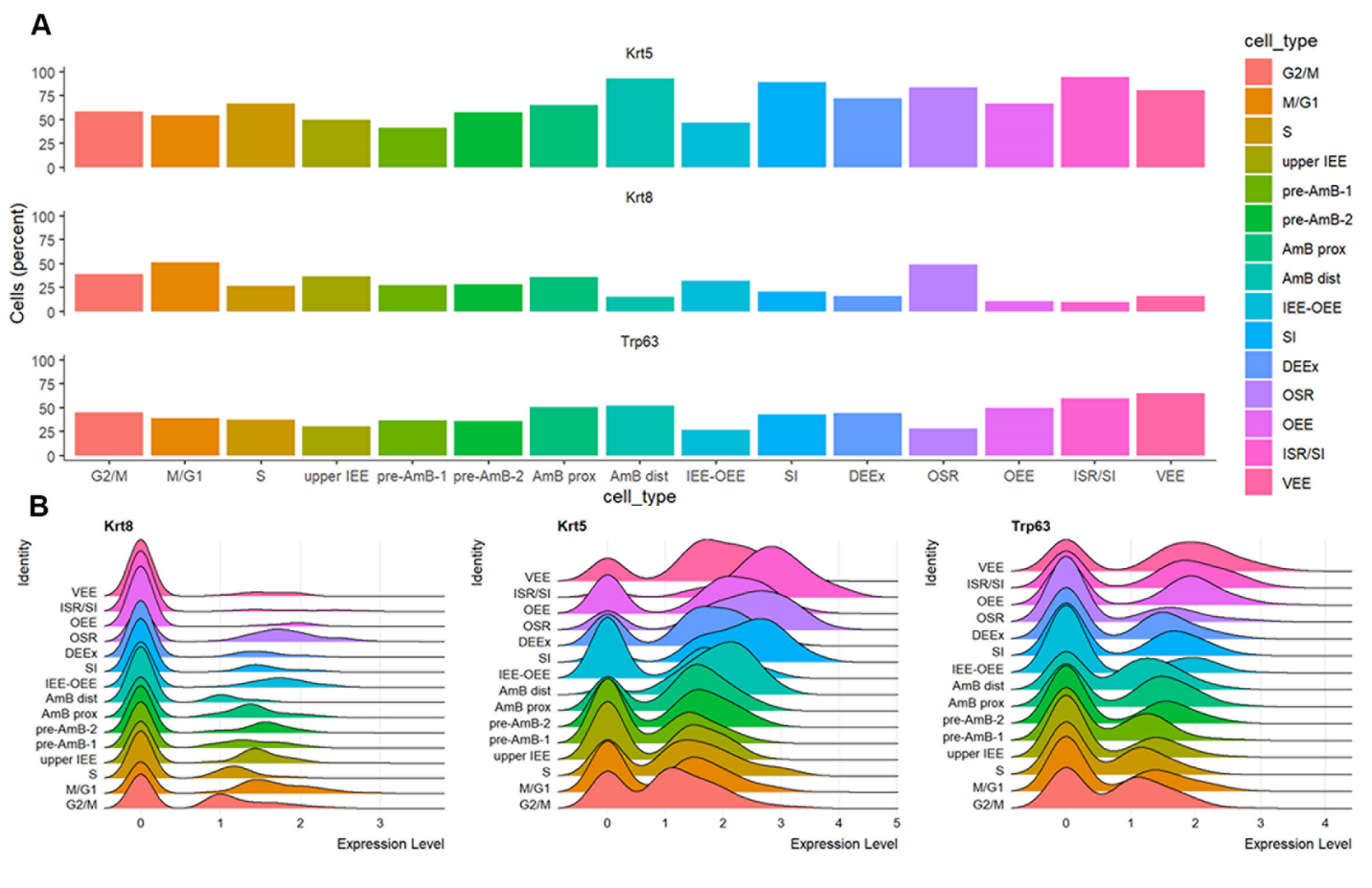

**Supplementary figure S2. *p63* expression coincides with Krt8-to-Krt5 transition during enamel organ development.** (**A, B**) The percentage of *p63*-expressing cells (A) and p63 expression level (B) are both positively correlated with the Krt8-to-Krt5 transition during enamel organ development.

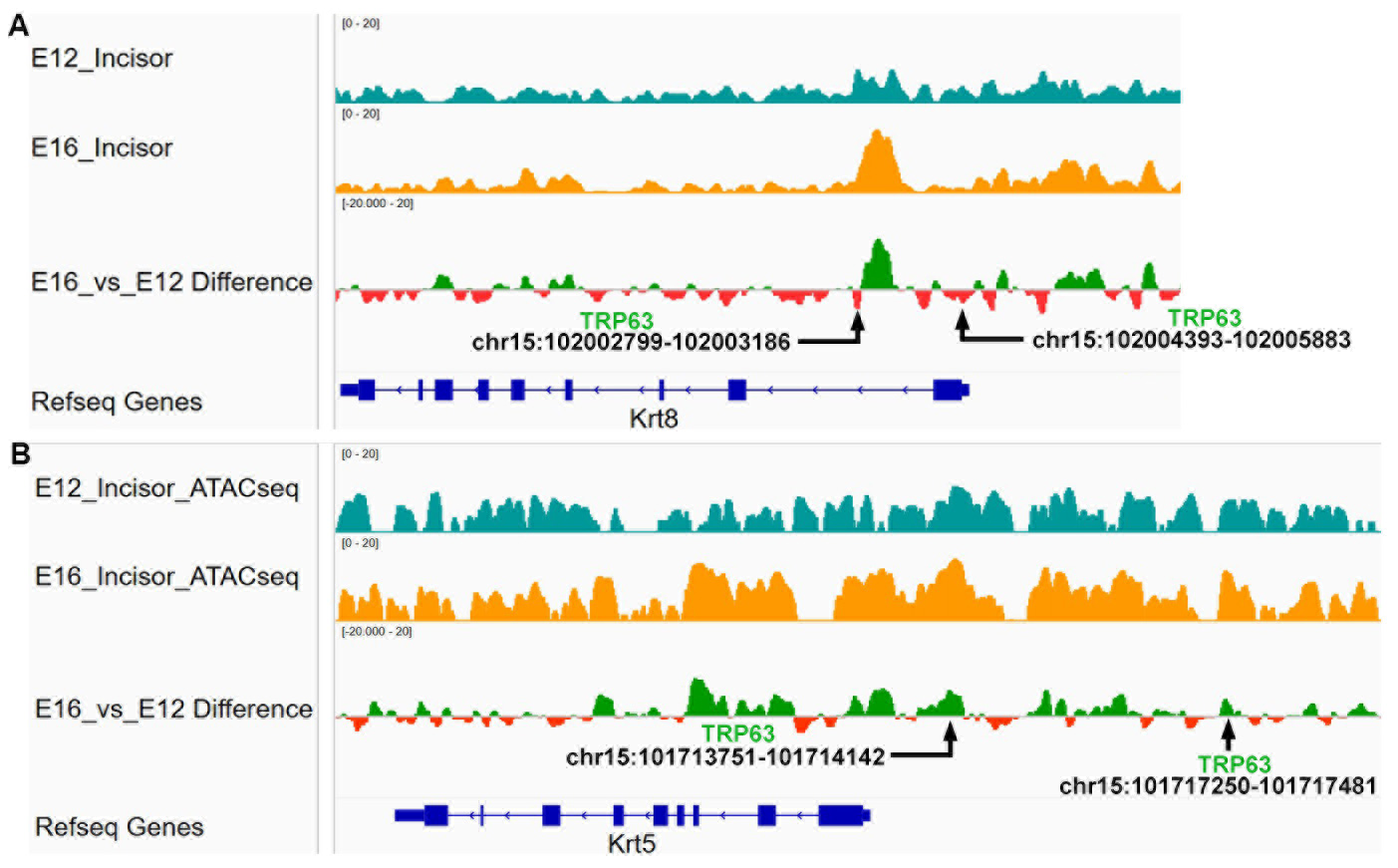

**Supplementary figure S3. Dynamic chromatin accessibility of *Krt8* and *Krt5* during mouse incisor development. (A, B)** ATAC-seq profiles demonstrating differential chromatin accessibility of *Krt8* (A) and *Krt5* (B) during mouse incisor development. The *Krt8* locus showed decreased chromatin accessibility at E16 compared to E12, whereas the *Krt5* locus exhibited increased chromatin accessibility. Red bars indicate regions of decreased chromatin accessibility at E16 relative to E12, while green bars highlight regions with increased chromatin accessibility. TRP63 binding sites with labeled chromosome coordinates indicate their presence in differentially accessible regions during the skin Krt8-to-Krt5 transition.
